# Genetic variation in aneuploidy prevalence and tolerance across the *Saccharomyces cerevisiae* phylogeny

**DOI:** 10.1101/2020.12.11.411785

**Authors:** Eduardo F.C. Scopel, James Hose, Douda Bensasson, Audrey P. Gasch

## Abstract

Individuals carrying an aberrant number of chromosomes can vary widely in their expression of aneuploidy phenotypes. A major unanswered question is the degree to which an individual’s genetic makeup influences its tolerance of karyotypic imbalance. Here we took a population genetics perspective to investigate the selective forces influencing aneuploidy prevalence in *Saccharomyces cerevisiae* populations as a model for eukaryotic biology. We analyzed genotypic and phenotypic variation recently published for over 1,000 *S. cerevisiae* strains spanning dozens of genetically defined clades and ecological associations. Our results show that the prevalence of chromosome gain and loss varies by clade and can be better explained by differences in genetic background than ecology. The phylogenetic context of lineages showing high aneuploidy rates suggests that increased aneuploidy frequency arose multiple times in *S. cerevisiae* evolution. Separate from aneuploidy frequency, analyzing growth phenotypes reveals that some backgrounds – such as European Wine strains – show fitness costs upon chromosome duplication, whereas other clades with high aneuploidy rates show little evidence of major deleterious effects. Our analysis confirms that chromosome amplification can produce phenotypic benefits that can influence evolutionary trajectories. These results have important implications for understanding genetic variation in aneuploidy prevalence in health, disease, and evolution.

**ARTICLE SUMMARY:** Aneuploidy, an imbalance in the normal chromosome copy number, is detrimental during human development; yet individuals show substantial variability in their aneuploidy susceptibility, suggesting the influence of genetic background on aneuploidy tolerance. Scopel *et al.* employed a population genetic approach to address this question, analyzing over 1,000 published *Saccharomyces cerevisiae* genomes. The results demonstrate that genetic background has a substantial effect on aneuploidy frequency and cellular tolerance of aneuploidy stress, presenting important new information on the forces that contribute to aneuploidy prevalence.

## INTRODUCTION

Chromosomal aneuploidy, in which cells carry too many or too few of individual chromosomes, can arise when chromosomes fail to segregate properly in meiosis or mitosis. An imbalanced karyotype is clearly detrimental during mammalian development, since amplification of all but a few specific chromosomes is inviable (MacLennan *et al.* 2015). Why aneuploidy is toxic remains an area of active research but is thought to result from imbalanced expression from affected chromosomes relative to the remaining genome (Pavelka and Rancati 2013; Oromendia and Amon 2014; Donnelly and Storchova 2015). Such imbalance could lead to myriad downstream problems, including proteostasis stress that may emerge when mismatched stoichiometries lead to protein misfolding and aggregation (Oromendia *et al.* 2012; Oromendia and Amon 2014; Donnelly and Storchova 2015). Despite the detrimental effects of aneuploidy during development, aneuploidy is common in other settings, notably cancers. Most solid tumors are aneuploid, yet cancer cells tolerate karyotype imbalance and in some cases thrive because of it (Holland and Cleveland 2012; Targa and Rancati 2018). In fact, chromosome amplification that emerges from extreme fitness pressure can present phenotypic variability and immediate phenotypic gains (Pavelka *et al.* 2010; Chen *et al.* 2012; Beach *et al.* 2017). The rapid onset of phenotypic differences plays an important role in selection: aneuploidy is frequently observed in drug-resistant *Candida albicans* and *Cryptococcus neoformans* pathogens that emerge from drug treatments, and microbes readily acquire chromosomes under extreme laboratory selective pressure (Hughes *et al.* 2000; Selmecki *et al.* 2006; Rancati *et al.* 2008; Selmecki *et al.* 2009; Yona *et al.* 2012; Filteau *et al.* 2015; Sunshine *et al.* 2015; Beaupere *et al.* 2018; Gilchrist and Stelkens 2019; Yang *et al.* 2019; Todd and Selmecki 2020). Why aneuploidy is toxic in some situations yet common in others is a matter of active debate but may reflect mechanistic differences in how cells handle the stress of aneuploidy depending on developmental stage, cell type, or species.

An understudied consideration is the influence of genetic background on aneuploidy tolerance, even within the same species. Genetic differences in aneuploidy tolerance have been suggested in several systems, but its influence is perhaps most clear in humans and mouse models of Down syndrome (DS). DS, caused by trisomy of human chromosome 21, is associated with a wide range of phenotypes; however, many of these macro and molecular phenotypes vary considerably across individuals (Antonarakis and Epstein 2006). For example, while DS is associated with a 50 fold increase in congenital heart defects, only half of DS patients suffer from heart problems (Li *et al.* 2016). Transcriptomic and proteomic analysis of patient-derived cell lines showed considerable variation in expression across individuals (Prandini *et al.* 2007; Liu *et al.* 2017). Several functionally related mRNA/protein classes display universal responses to trisomy 21, whereas other functional processes vary across lines, revealing the genetic influence on cellular susceptibilities. Some of these differences may be encoded on the amplified chromosome, with profound influences on DS severity. A recent study of DS patient genomes identified a dearth of deleterious alleles on sampled chromosome 21, which was inferred to reflect survivor bias in tolerated variants (Popadin *et al.* 2018). However, the influence of broader genetic background effects has been less clear, as there has been no systematic study exploring the aneuploidy tolerance across genotypes.

We recently showed that genetic differences between laboratory and wild strains of budding yeast *Saccharomyces cerevisiae* influence the tolerance of chromosome amplification. Laboratory strain W303 is highly sensitive to amplification of any of the 16 yeast chromosomes, showing extreme growth defects, metabolic limitations, transcriptome reorganization, cell-cycle defects, and proteostasis stress including protein aggregation and difficulties degrading misfolded proteins (Torres *et al.* 2007; Torres *et al.* 2010; Oromendia *et al.* 2012; Sheltzer *et al.* 2012; Thorburn *et al.* 2013; Dephoure *et al.* 2014; Dodgson *et al.* 2016; Brennan *et al.* 2019). However, natural aneuploid yeast strains studied to date do not show these phenotypes, and instead grow with more mild growth reduction, no evidence of the stress response, and little evidence of metabolic or proteostasis stress (Hose *et al.* 2015; Gasch *et al.* 2016). We recently mapped the difference in aneuploidy tolerance between W303 and wild strains to RNA binding protein Ssd1, which is defective in W303 (Hose *et al.* 2020). Indeed, deletion of Ssd1 recapitulates aneuploid-W303 phenotypes in several genetic backgrounds with different chromosome amplifications, revealing a generalizable role for Ssd1 in tolerating chromosome duplication. Since then, work by Larrimore *et al.* showed that aneuploidy in a different laboratory strain derived from S288c, which expresses full-length Ssd1, produces no observable proteostasis defects reported to be a hallmark in aneuploid W303 (Larrimore *et al.* 2020).

A remaining question is whether there is natural variation, outside of aberrant laboratory strains, in aneuploidy tolerance. *S. cerevisiae* is an excellent model to explore this question. Several studies have documented aneuploidy in large-scale sequencing efforts, demonstrating that chromosome imbalance is not uncommon and can be found associated with several ecological settings, especially clinical isolates and industrial strains (Hose *et al.* 2015; Strope *et al.* 2015; VAN DEN Broek *et al.* 2015; Gallone *et al.* 2016; Zhu *et al.* 2016; Gorter de Vries *et al.* 2017; Duan *et al.* 2018; Fay *et al.* 2019). Peter *et al.* recently sequenced over 1,000 *S. cerevisiae* strains collected from diverse environments, including natural ecologies such as fruit and trees, yeast used in industries spanning beverage, bread, and biofuel production, and human isolates from infections and other human-associated environments (Peter *et al.* 2018). Their phylogenetic analysis identified at least 26 defined genetic clades associated with geography or ecology, along with many other “mosaic” strains that show recent genetic admixture. That study also generated growth phenotypes for those strains growing in different environments. Although not a focus of the study, a fifth of the sequenced strains harbor atypical aneuploid karyotypes. Here we explore patterns of aneuploidy and its phenotypic effects to understand how genotype and ecology influence aneuploidy prevalence and tolerance. Although intertwined with ecology, our results demonstrate that genetic background alone can predict differences in aneuploidy frequency and phenotypes. We present models for the evolutionary forces acting on aneuploidy in the species.

## METHODS

### Strain analysis

Genotypes, clade classifications, and phenotypes were taken from Peter *et al*. (2018). We confirmed all aneuploidy calls by comparing average read depth (calculated from 1-kb non-overlapping sliding windows) between chromosomes. In total, Peter *et al.* assigned clade classifications to 962 strains (which included 150 strains designated as ‘mosaic’) and ecological associations for an overlapping 984 strains. The European Wine clade and mosaic strains analyzed here included all subclades in each group identified by Peter *et al*. Natural strains in Fig 2 included flower, fruit, insect, nature, soil, and tree ecotypes, and industrial strains included bakery, beer, fermentation, palm wine, industrial, and bioethanol ecological groups.

Strains were partitioned based on aneuploidy calls if they displayed gain of Chromosome I (Chr 1) only, loss of one or more chromosomes bigger than Chr 1 (Chr 2-16) without any gains (“loss only”), gain of one or more chromosomes bigger than Chr 1 without any losses (“gain only”), and strains that had gained and lost different chromosomes bigger than Chr 1 (“mixed gain and loss”). Enrichment for strains with aneuploidy was assessed by comparing to the total set of strains using Fisher’s exact test and Benjamini and Hochberg false discovery rate (FDR) correction, taking FDR < 0.05 as significant and FDRs that just missed the cutoff (as specified in each figure legend) as marginally enriched. Only groups of at least 8 strains are represented in Fig 2.

### Verification of aneuploidy calls, heterozygosity estimation and phylogenetic tree construction

Reads were trimmed for all sequences from Peter *et al.* (2018) using Trimmomatic (version 0.33; Bolger *et al.* 2014) with default settings and read quality, and the efficacy of the trimming was assessed with FastQC (version 0.11.8; http://www.bioinformatics.babraham.ac.uk/projects/fastqc/). Trimmed reads were mapped to the *S. cerevisiae* reference genome (SacCer_Apr2011/sacCer3 from UCSC) with Burrows-Wheeler Aligner (bwa mem, version 0.7.17; Li and Durbin 2009). A consensus sequence was generated for each strain using SAMtools mpileup (version 1.6; Li *et al.* 2009) and BCFtools call -c (version 1.6; Li *et al.* 2009). Indels were removed in the mpileup step (option -I) and a maximum read depth of 100,000 reads was allowed. Levels of heterozygosity were estimated from the resultant variant call format (vcf) files using vcf2allelePlot.pl with default parameters (Bensasson *et al.* 2019). Consensus genome sequences in fasta format were extracted from the vcf file of each strain using seqtk (available at https://github.com/lh3/seqtk), and for each strain consensus bases with a phred-scaled quality score below 40 were counted as missing data (converted to N). Because indels were removed when mapping reads to the reference, all genome sequences are already mapped to the same coordinates as the reference sequence and there was therefore no need for a multiple alignment step. For tree construction, the alignments of all sixteen chromosomes were concatenated into a single 12,071,326 bp genome-wide alignment using alcat.pl (Bensasson *et al.* 2019), with 1,100,664 variable sites. From this alignment, a maximum-likelihood tree was inferred using IQ-TREE (Version 1.6.5; Nguyen *et al.* 2018) with a general time reversible evolutionary model with unequal rates and unequal base frequencies, a discrete γ-distribution to estimate heterogeneity across sites (GTR+G), and 100 nonparametric bootstrap replicates.

*SSD1* forward DNA coding sequence was extracted from the fasta file for each strain using faChooseSubseq.pl with the -r option to recover the correct strand (Bensasson *et al.* 2019). Aligned sequences revealed 282 variant sites. RAxML (version 8.2.11 (Stamatakis 2014)) was used to infer a maximum-likelihood tree with a general time reversible evolutionary model, a discrete γ-distribution to estimate heterogeneity across sites (GTRGAMMA), and 100 bootstrap replicates (seed -x 12345). Similar methods were applied to generate the Ssd1 protein tree. The data table and phylogenetic trees generated are available at https://github.com/bensassonlab/data/tree/master/scopel_etal20/.

### Logistic regression analysis of chromosome gain

To directly compare clade, ecological origin, ploidy and heterozygosity levels as predictors of the probability of Chr 2-16 gain, we considered them together in logistic regression models. Out of the 812 strains with aneuploidy information and assigned to clades by Peter *et al*, we focused these logistic regression analyses on 621 strains that were euploid (N=520) or aneuploids that had gained chromosomes bigger than Chr 1 without any additional chromosome losses (N=101). This filtered dataset excludes (i) 86 strains that had been manipulated (*i.e.* in which the HO locus was deleted), (ii) 45 strains that were monosporic derivatives (Strope *et al.* 2015) and therefore less likely to be aneuploid (Zhu *et al.* 2016), (iii) 6 strains that were haploid and therefore effectively had missing data for heterozygosity, (iv) 2 strains (CBS382 and CBS1593) that our phylogenetic analysis showed were not in the clades assigned by Peter *et al.* (2018).

To identify the strongest predictors of the frequency of Chr 2-16 gain by logistic regression in R, we used the generalized linear model (glm) function with binomial errors. More specifically, the response variable was set as Chr 2-16 gain (CG, true or false) and the initial model included 4 explanatory variables: clades (C, 26 levels), ecological origin (E, 23 levels from Peter *et al*), ploidy (P, diploid, polyploid) and heterozygosity (H, continuous). We simplified a maximal model (CG ~ C*E*P*H) to minimal adequate models by applying chi-squared deletion tests, as recommended by (Crawley 2013) for analysis of data where variables are correlated. We ensured that all conclusions are not affected by the order of factor deletion and also compared the Akaike Information Criterion (AIC) estimator for our models using the summary glm function. The fullest models would not converge with all explanatory variables in a single model together with all their possible interactions. Instead, for the maximal model, we considered these 4 variables together with as many of their two-way interactions as possible (CG ~ C + E + P + H + C:H + E:P + E:H + P:H); all except clade:ecology (C:E) and clade:ploidy (C:P) were included. None of the two-way interaction terms were statistically significant (*P* > 0.05) and are therefore not discussed in the Results. The R script and data table used for the logistic regression along with files for other analyses presented here are available at https://github.com/bensassonlab/data/tree/master/scopel_etal20.

The potential non-additive effects of variables that are described by statistical interaction terms can be important for model interpretation. Because maximal models did not converge when three-way interaction terms were included above, we also predicted the gain of Chr 2-16 (true or false) response from two additional models each with 3 explanatory variables: one for ecology, heterozygosity and ploidy along with all possible interactions (CG ~ E*H*P), and a second model for clade, heterozygosity and ploidy with all possible interactions (CG ~ C*H*P). We again simplified the models using the standard approach of chi-square tests to compare nested models with the anova function in R (Crawley 2013). No two-way or three-way interactions were statistically significant (*P* > 0.05) and both models simplified the same way as in the logistic regression analysis described above.

To estimate the number of times that aneuploidy frequency has changed during the evolution of *S. cerevisiae*, we simplified the 26 clades defined by Peter *et al* (2018) by *a priori* contrasts (Crawley 2013). More specifically, we used nested models and chi-square tests with the anova function (Crawley 2013), and the *S. cerevisiae* phylogeny (Fig 6A) to decide which *a priori* contrasts to use. We used the phylogeny by comparing each clade to its most closely-related clade and merged clades if there was no significant difference between component clades in the probability of Chr 2-16 gain, using a significance threshold of *P*=0.05. Once all sister clades with similar aneuploidy frequency were merged, we tested whether our model was significantly worse when we replaced all clades that appeared to have an ancestral rate of Chr 2-16 gain (clades that are not highlighted in Fig. 6A) with a single group. Maximum likelihood estimates of the mean probability of Chr 2-16 gain for each group were extracted from the minimal adequate model using the predict function in R.

We controlled for the possibility that recent common ancestry of Chr 2-16 gains could inflate aneuploidy rates for some clades, by re-running the logistic regression analysis after excluding close relatives. To do this, we used the dnadist function of PHYLIP to estimate pairwise genetic distances for all 621 strains (version 3.697 (Felsenstein 1989)). Using a python script (removeSimilarStrains.py; https://github.com/bensassonlab/data/tree/master/scopel_etal20), we randomly chose a single strain from each group of very similar strains (pairwise genetic distance < 0.000007). Phylogenetic analysis of the 453 strains that remained indicated successful removal of closely related strains with identical aneuploid karyotypes. In contrast, many of the remaining 31 polyploids clearly share a common polyploid ancestor (*e.g.* African Beer polyploids in Fig S1B). Logistic regression analysis of genetic clade, ecology, heterozygosity and ploidy showed statistically significant two-way interactions involving ploidy (*P* = 0.01 to *P* = 0.05), suggesting that polyploidy might not increase aneuploidy rates in all genetic backgrounds or environments. To test whether changes in aneuploidy prevalence arose multiple times, we therefore excluded polyploids and repeated the logistic regression analysis on Chr 2-16 gain in 422 diploids, this time using the phylogeny that excludes close relatives (Fig S1B) to determine *a priori* contrasts.

### Phenotype analysis

Phenotype scores included colony size after growth for defined periods in different environments (Peter *et al.* 2018) and sporulation measurements from (De Chiara *et al.* 2020). Differences in colony sizes between aneuploid and euploid groups were scored by a Wilcoxon rank-sum test (implemented in R version 3.5.1 with a continuity correction) taking FDR < 0.05 as significant. To ensure that the European Wine sensitivity was not due to an unusually high aneuploidy burden in these sampled strains, we removed 4 outlier strains whose total genome content was greater than the largest sake genome (27.3 Mb), which is the aneuploid group with among the tightest distribution of genome contents. The median genome size of the remaining European Wine strains was not different from French Dairy, Sake, and Mosaic aneuploids; yet these European Wine aneuploids grew slower than euploids (*P* = 0.02) and showed fewer asci and more dyads (*P* <0.01, one-sided Wilcoxon test). Where indicated, colony-size scores from cells grown in stress conditions were normalized to that strain’s colony-size score from rich YPD media, thereby representing the relative change in colony size in each stress. To score phenotypic benefits afforded by amplification of specific chromosomes (Fig 5B), FDR was calculated using only tests where the median aneuploid growth rate was equal to or better than the euploid growth rate.

### Ssd1 variation

To test if Ssd1 B variants were enriched among aneuploid strains, beyond what is expected by skews in the genetic makeup of this group, we did the following: Among aneuploid strains that had amplified Chr 2-16 (without any losses), we calculated the number of strains from each of the Peter *et al.* clades. We estimated clade-specific AA, AB, and BB genotype frequencies, based only on euploid strains in the datasets, and then used these frequencies to estimate the expected frequencies of the aneuploid group based on its clade proportions. Frequencies of the genotypes were not different from expectation (*P* = 0.98, Chi-squared test).

### Experimental methods

The *SSD1* coding sequence plus 1,000 bp upstream and 337 bp downstream was cloned from YPS1009 or from S288c strain BY4741 into a low-copy CEN plasmid for complementation as previously described (Hose *et al.* 2020). Site-directed mutagenesis of the *SSD1^YPS1009^* expressing plasmid was used to introduce the S1190G and A1196P substitutions into a YPS1009 backbone. All plasmids were sequence verified. Plasmids were transformed into wild-type and *ssd1D* aneuploids of the haploid YPS1009_Chr12 and NCYC110_Chr8 backgrounds (each disomic for the affected chromosome), aneuploidy was verified by qPCR as previously described (Hose *et al.* 2020), and growth rates in rich medium were measured in biological triplicate. Data in Figure 6C show the relative growth rates of *ssd1D* aneuploid strains transformed with each plasmid relative to the wild-type aneuploid transformed with the empty vector.

Data availability statement: Strains and plasmids are available upon request. Data files and scripts are available at https://github.com/bensassonlab/data/tree/master/scopel_etal20/. The authors affirm that all data necessary for confirming the conclusions of the article are present within the article, figures, and tables or previously published.

## RESULTS

We began by investigating the levels and types of chromosomal aneuploidies in 1,011 strains sequenced by Peter *et al*. (2018). Of the 217 (21%) aneuploid strains, 26 were reported to show only chromosomal loss, whereas 191 strains amplified one or more of the 16 yeast chromosomes, a third of which duplicated two or more chromosomes. Chromosome I (Chr 1), both the smallest and carrying the fewest genes, was the most frequently duplicated, in 60 (27.6%) aneuploid strains. It was also the most frequently lost (14/217, 6.5%). The higher frequency could be due to increased rate of segregation errors; however, given its small size and low gene count this chromosome is least likely to incur large fitness defects when amplified. Consistent with this notion, we confirmed the result of Peter *et al.* that the frequency of chromosome amplification is negatively correlated with chromosome size and the covarying gene content (R^2^ = 0.38, Fig 1A), and in general smaller chromosomes were more often duplicated than larger chromosomes. This trend is reminiscence of the size-specific differences in fitness costs of chromosome amplification, measured across many backgrounds (Hose *et al.* 2015) and in a single laboratory strain in which most of the individual chromosomes have been duplicated (Torres *et al.* 2007; Gilchrist and Stelkens 2019). Interestingly, there was an even stronger negative correlation for the rate of chromosome loss and chromosome gene content (R^2^ = 0.53, Fig 1B). Together, these results are consistent with the model that substantial alteration in chromosome/gene content is generally selected against in *Saccharomyces cerevisiae*, with smaller chromosomes likely to incur smaller fitness costs, contributing to their higher rate of detection in yeast populations. However, the relationship was not perfect. The second most commonly amplified chromosome in the 1,011 strains is Chr 9 (53 strains), the 4^th^ smallest chromosome previously observed to amplify frequently in clinical isolates (Zhu *et al.* 2016), followed by Chr 8, 11, and 3 (24-28 strains each); although the frequency of aneuploidy did not correlate perfectly with size, all of these chromosomes are smaller than the median chromosome length.

**Figure 1.**
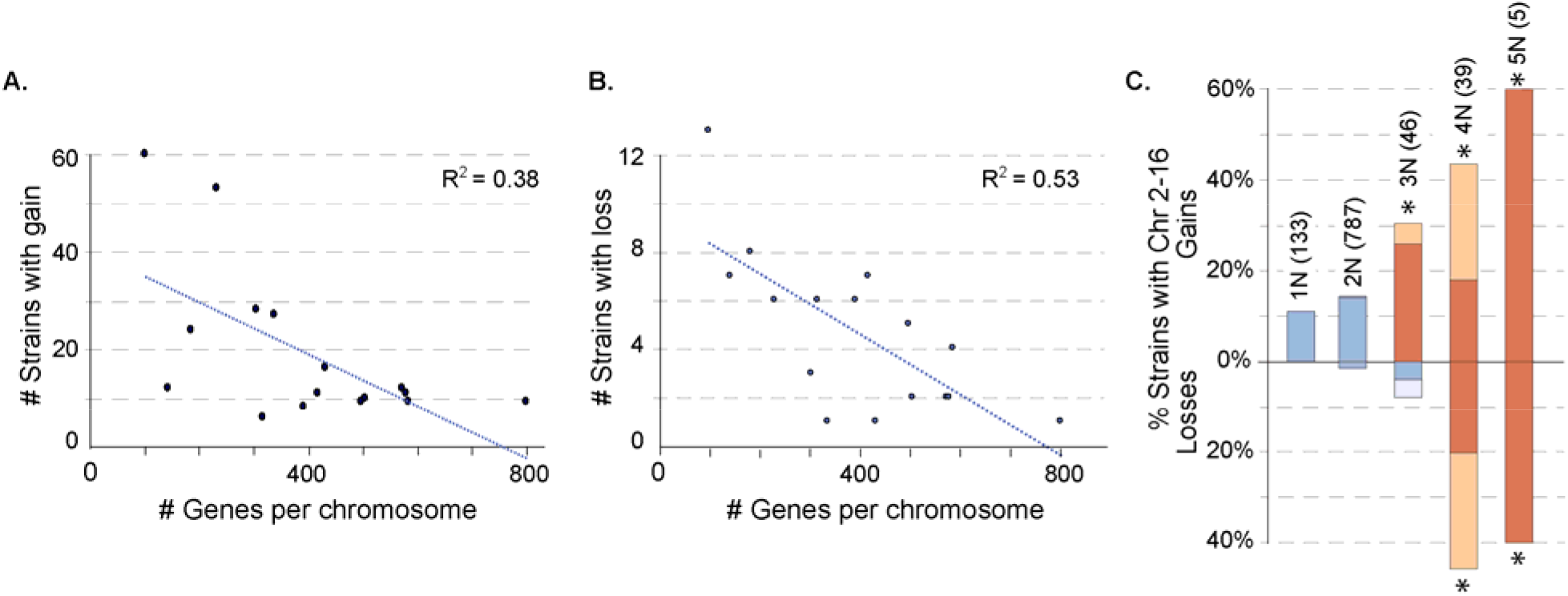
Frequency of gain and loss correlates with chromosome gene content. The number of strains that had gained (A) or lost (B) each chromosome (regardless of other chromosome patterns in that strain) is plotted against number of genes on that chromosome. The percent variance explained is shows as the R^2^ of the linear fit. Chromosome size (which covaries with gene content) is depicted in Figure 3. (C) The proportion of strains from each group that had gained or lost chromosomes larger than Chr 1 (Chr 2-16). Dark colored bars indicate strains with only chromosome gain or loss; light colored bars indicates strains with both gain and loss and were counted in both the gain and loss categories. The number of strains per ploidy group is listed in parentheses. Frequencies beyond chance were based on total values above or below the line and are highlighted in orange (asterisk, FDR < 0.05).

Whole-genome duplication precedes chromosome instability in cancer cells, giving rise to highly aberrant karyotypes (Storchova and Pellman 2004; Storchova 2014). The same forces may occur in yeast. We expanded on previous observations that aneuploidy is more common in polyploids (Mayer and Aguilera 1990; Storchova 2014; Zhu *et al.* 2016; Duan *et al.* 2018; Gilchrist and Stelkens 2019) with additional resolution. Because amplifications and losses may be subject to different selective pressures, we analyzed separately strains that showed only chromosome loss, those that gained only the smallest Chr 1, and strains that amplified chromosomes larger than Chr 1 (Chr 2-16), with or without other chromosome losses. As ploidy increases, aneuploidy frequency increases, for both chromosome gains and losses (Fig 1C). Interestingly, haploid gains of Chr 2-16 (in the absence of any chromosome loss) were only slightly less prevalent (11%) than in diploids (14%; Fisher’s exact test, *P*=0.4), but triploids had over double the frequency of Chr 2-16 gain (30%, Fisher’s exact test, *P=*0.005). The frequency of chromosome loss (exclusive of or including other gains) is increasingly prevalent as ploidy increases, and aneuploidy becomes especially common in tetraploids and pentaploids (Fig 1C). This may arise from a higher rate of chromosome slippage as ploidy increases (Storchova *et al.* 2006; Marco *et al.* 2013), but it could also be more tolerated if the load of genome imbalance of single-chromosome alteration compared to the haploid genome content is smaller.

We also noticed that aneuploid strains were more heterozygous than others. Even when controlling for ploidy effects by examining only diploids, cells with any type of aneuploidy were more heterozygous (median 0.05%, N=145) than euploids (median 0.04%, N=642; Wilcoxon test, *P=*0.0002). More specifically, heterozygosity was elevated for strains that exclusively gained Chr 2-16 (median 0.05%, N=108; Wilcoxon test, *P* = 0.0001), although the higher heterozygosity was not significant for the few strains that exclusively lost Chr 2-16 (median 0.08%, N=10; Wilcoxon test, *P =* 0.3). In subsequent analyses, we therefore consider the effects of ploidy and heterozygosity alongside those of genetic lineage and ecological origin.

### Aneuploidy frequency varies across lineages and ecological groups

Peter *et al.* identified 26 distinct genetic lineages or clades, which we recapitulated in our own phylogenetic analysis (see Fig 6A), along with many more mosaic strains showing recent genetic admixture between these groups. One question is if different lineages show different propensities for aneuploidy. We therefore scored the frequency of aneuploidy by genetic lineage as defined by Peter *et al*, again distinguishing between gains only, losses only, or gains with losses of Chr 2-16 (Fig 2). As previously noted (Gallone *et al.* 2016; Peter *et al.* 2018; Fay *et al.* 2019), strains in the Ale, Sake, and Mixed Origin lineages display high rates of chromosome amplification, with 40-50% of individuals carrying extra copies of chromosomes larger than Chr 1 (p < 0.008, Fisher’s exact test). In contrast, the European Wine clade was significantly under-represented for strains with extra chromosomes. In addition to gains, African Beer, Ale and Mixed Origin clades also showed a prevalence for chromosome loss, in the context of other amplifications (p < 4e-7, Fisher’s exact test) or in strains that showed only loss (Fig 2). All three clades are also enriched for polyploids.

**Figure 2.**
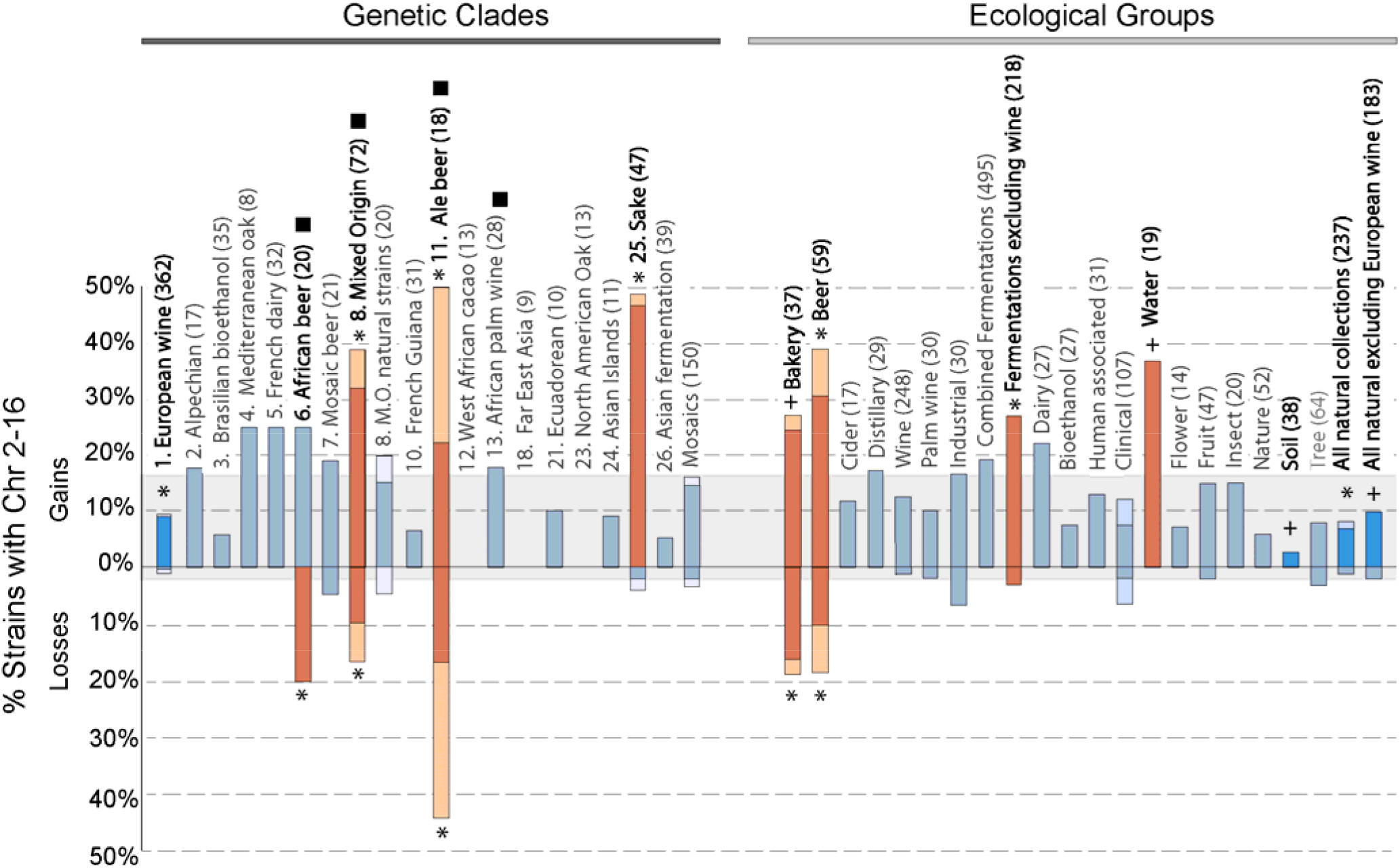
Aneuploidy prevalence varies by clade and ecological group. As shown in Fig 1C. Groups significantly enriched (orange) or depleted (sky blue) of aneuploid isolates compared to the frequency across all strains (grey box) are highlighted, and the number of strains in each group indicated in parentheses. Asterisk indicates FDR < 0.05 for total gains or losses, and plus indicates p<0.05 and FDR < 0.13. Black square indicates clades enriched for polyploids (FDR < 0.05). All groups with at least 8 strains are shown.

There were also several striking differences when strains were grouped instead by ecology. As expected based on previous studies (Gallone *et al.* 2016; Gorter de Vries *et al.* 2017; Kadowaki *et al.* 2017; Peter *et al.* 2018), strains used in some industries (including beer, bread, and sake but not wine production) showed elevated rates of chromosome amplification. In contrast, strains isolated from nature, notably those from soil or all natural ecologies combined (Fig 2), were significantly depleted of aneuploids compared to the total set of 1,011 strains. However, a major challenge with this interpretation is that some ecologies are tightly associated with genetic lineage. For example, sake-making strains form a unique genetic clade, as do ale strains that were recently shown to be hybrids between Asian and European wine strains (Fay *et al.* 2019). In this dataset, natural strains are heavily enriched for the European Wine lineage, itself underrepresented for aneuploids. Understanding how environment affects aneuploidy prevalence thus requires methods that disentangle genotype from ecology.

### Genetic background alone can predict differences in aneuploidy prevalence

Differences in aneuploidy prevalence could result from the direct action of ecological selection, or it could arise due to inherent differences in aneuploidy tolerance according to genetic background. If ecology directly affects the prevalence of aneuploids, then it should improve predictions of aneuploidy rates when considered alongside other factors. Such an analysis is complicated by the co-variation of ploidy, heterozygosity, clade, and ecology, which confounds the forces driving aneuploidy frequency. For example, strains from the Mixed Origin clade are more likely to be aneuploid (Fisher’s exact test, *P* = 1×10^−9^), but they are also more often polyploid (*P* = 6×10^−14^, Fisher’s exact test) and more heterozygous (*P* < 2×10^−16^, Wilcoxon test) than other clades. Furthermore, many of these are bakery or clinical strains for which there is a known association between ploidy and aneuploidy (Zhu *et al.* 2016).

To begin to disentangle factors that influence aneuploidy prevalence, we performed a logistic regression that included clade designation, ecological source, ploidy, and heterozygosity to predict the frequency of Chr 2-16 gain (in the absence of chromosome loss for clarity of interpretation). When simplifying a maximal model that included all variables (clade, ecology, ploidy, heterozygosity) and most two-way interactions (except clade:ecology and clade:ploidy, see Methods), we found no effect of ploidy beyond what was explained by other factors (d.f.=1, *P*>0.1). Because clade and ecology covary, model simplification produced two different minimal adequate models depending on the order of deletion. These two models were not significantly worse than any more complicated models (see Methods). (i) The first model (CG ~ C; AIC=504.54) predicted differences in Chr 2-16 gain using only genetic background, and the model revealed major differences among genetic clades (17.9% of deviance, df = 25, *P* = 9×10^−11^). (ii) The second model (CG ~ E + H, AIC = 499.25) predicted the probability of aneuploidy from ecological source (14.8% of deviance, df = 22, *P* = 1×10^−8^) and heterozygosity (1.8% of deviance, df=1, *P*=0.002). Clade explains a larger fraction of the differences (deviance) in chromosome gain in the first model than ecology does in the second model. Furthermore, the second model must also include heterozygosity for explanatory power, which covaries with genetic clade and may act with ecology as a surrogate for clade. Together, these data suggest genetic clade is a better predictor of the differences in chromosome gain than ecology (see Discussion).

### Increased aneuploidy frequency arose multiple times in S. cerevisiae evolution

If genetic clade is the best predictor of aneuploidy prevalence, then could genetic differences in aneuploidy tolerance have arisen in the evolution of the species, and could such changes have arisen multiple times? To address these questions, we further simplified the logistic regression model from 26 clades to larger groups defined using phylogenetic relationships and aneuploidy frequency (see Fig 6A and Methods). For example, the Alpechin clade is most closely-related to the European Wine clade (Fig 6A) and also shows a similar prevalence of chromosome gain (d.f. *=* 1, *P =* 0.3); these clades can therefore be combined into a larger clade with a single shared probability of aneuploidy. In contrast, Sake and Asian Fermentation strains are sister clades but differ in the prevalence of chromosome gain (d.f. *=* 1, *P =* 6×10^−7^), so the propensity for aneuploidy must have changed during the divergence of these lineages. In this way, simplifying the clade categories on the basis of their phylogenetic relationships (Fig 6A) revealed that the probability (frequency) of Chr 2-16 gain (P_Chr2-16_) increased through multiple events: along the lineages leading to (i) Sake (P_Chr2-16_ = 0.59), (ii) French Dairy and African Beer (P_Chr2-16_ = 0.27), (iii) Ale beer and Mixed Origin clades (P_Chr2-16_ = 0.39 for the reduced set of strains analyzed here, see Methods), and (iv) Mosaic Beer (P_Chr2-16_ = 0.38). In this simplest model, the clade variable simplified to 5 groups: the 4 high-aneuploidy groups capturing 6 defined genetic clades and a fifth group where the same ancestral aneuploidy probability was estimated for all clades (“ancestral rate” strains, P_Chr2-16_ = 0.09). This minimal adequate model with only 5 parameters (AIC= 480.33) predicts differences in the probability of Chr 2-16 gain almost as well (14.7% of deviance) as the model with 26 clades (17.9% of deviance, AIC=504.54) without being significantly worse (−3.2% of deviance, d.f. *=* 21, *P =* 0.7). These results suggest that the ancestral state is a lower frequency of aneuploidy at ~9%, similar to that seen for the European Wine lineage, but that multiple independent events along the phylogeny have resulted in higher frequencies of chromosome gain (see Discussion). This result is not an artefact of clonal expansion of aneuploid strains: we repeated our analysis after removing strains that were closely related to others in the dataset (see Methods), which confirmed evidence for multiple independent increases in the prevalence of Chr 2-16 gains across the *S. cerevisiae* phylogeny (Fig S1).

### Forces influencing aneuploidy occurrence

The differences in aneuploidy prevalence across clades could emerge for several reasons. On the one hand, amplification of specific chromosomes has been proposed to benefit industrial strains, and aneuploidy could have been selected for during domestication of niche-associated clades. On the other hand, selective pressures may vary considerably in different ecologies (including those tightly associated with clades), such that relaxed purifying selection explains differences in prevalence. Growth, meiosis, and competition in mixed communities are likely less important in managed industrial processes, and thus selection against detrimental chromosome duplications may be relaxed. But a third possibility is that different genetic lineages may have acquired inherently different abilities to tolerate chromosome amplification, such that fitness costs are different in different lineages. Below we combine genetic and phenotypic analysis to address these possibilities.

### No evidence for selection of specific chromosome amplifications in industrial strains

As highlighted above, ale and sake producing strains as well as those linked to baking and beer production are enriched for chromosome amplification. It has been proposed that amplification of specific chromosomes has been selected for, presumably due to up-regulated production of genes enhancing desirable traits (Gorter de Vries *et al.* 2017; Kadowaki *et al.* 2017). This model posits that strains associated with different industries should be enriched for specific aneuploid karyotypes. To test this, we scored the frequency of each chromosome amplification compared to what was expected by chance (based on the total set of observed aneuploids in this dataset). In most cases we found that lineages with a high propensity for chromosomal aneuploidy did not share amplification of the same chromosomes (Fig 3). For example, while over 50% of strains in the Ale lineage carried extra chromosomes, there was no enrichment for any single chromosome in the group. Ale strains were also enriched for chromosome loss compared to other strains (*P* = 1×10^−10^, Fisher’s exact test); however, which chromosome(s) were lost also varied by strain.

**Figure 3.**
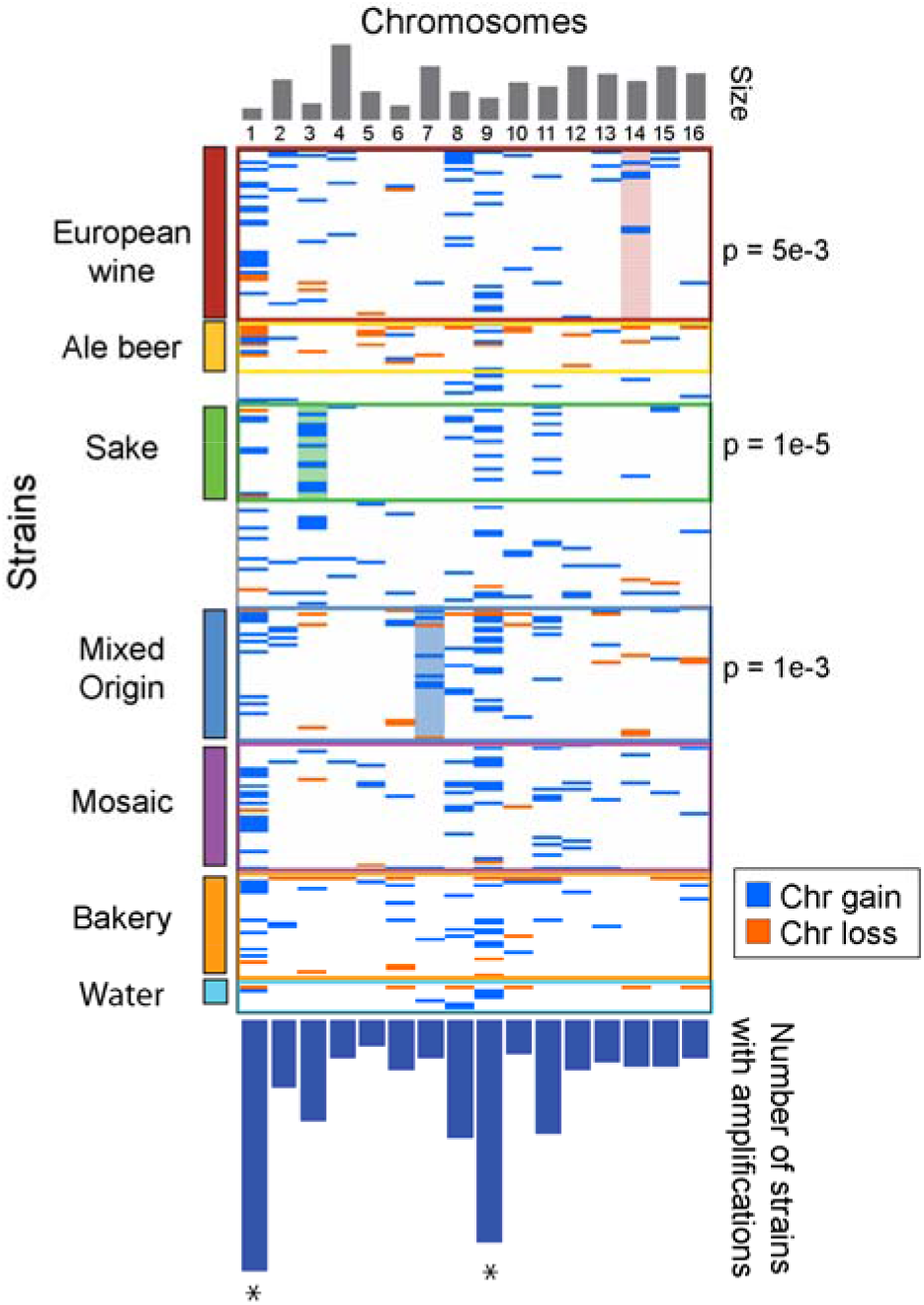
Most clades are not associated with specific karyotypes. The figure shows gain (blue lines) or loss (orange lines) of each chromosome (columns) in aneuploid strains (rows) belonging to each clade or select ecology groups. Not all ecologies are shown in the figure. Clades with significant enrichment for particular chromosomes are highlighted with p-value of the enrichment shown to the right. The top panel represents chromosome size and the bottom panel represents the fraction of all strains in the analysis in which each chromosome was amplified.

Somewhat surprisingly, a similar situation is seen for sake-producing strains, one of the few cases in which amplification of a specific chromosome is known to contribute desirable flavor characteristics. High pyruvate levels lead to off flavors in sake, and thus low pyruvate production is thought to have been selected for in modern-day sake-producing strains (Horie *et al.* 2010; Agrimi *et al.* 2014). Kadowaki *et al.* showed that sake strain TCR7 carrying an extra Chr 11 out-performed euploid variants of the same genotype (Kadowaki *et al.* 2017). We found that while a fifth of the aneuploid sake strains sequenced by Peter *et al.* carried extra Chr 11, there was no enrichment above expectation, based on the distribution of Chr 11 amplification across all strains. Instead, we found enrichment for Chr 3 duplication, which was often found with other aneuploidies (Fig 3). Removing closely related strains (see Fig S1) or simply mapping karyotypes onto the phylogeny (Fig S2) shows that Chr 3 and 11 amplifications are not due to clonal expansion of the same strain. These results disfavor the model that high aneuploidy levels are due to selection for specific chromosome amplifications. Instead, it suggests that Sake strains have a higher rate of segregation errors, a higher tolerance of karyotype imbalance, or both (see below and Discussion).

### Phenotypic variation points to underlying differences in aneuploidy tolerance

Wild aneuploids studied to date are not nearly as sensitive to chromosome amplification as laboratory strain W303, based on growth rate differences compared to their euploid cousins (Hose *et al.* 2015; Gasch *et al.* 2016; Hose *et al.* 2020). Nonetheless, a remaining explanation for differences in aneuploidy frequency is that wild strains still vary (albeit less so than compared to W303) in their ability to handle extra chromosomes. We reasoned that growth phenotypes of euploid and aneuploid strains within each clade may provide clues. Peter *et al.* measured colony size as a proxy for growth rate for 1,011 strains grown in 36 conditions, including growth on rich medium without added drugs or stresses. While they reported a slight but statistically significant fitness defect across aneuploids pooled across all conditions (and regardless of gain/loss type), we sought to look in more detail. We again considered separately strains with only chromosome loss, only Chr 1 amplification, or amplification of Chr 2-16 (here, regardless of other losses). We started by analyzing growth in rich medium, which is among the most optimal conditions for yeast growth, to assess if observed aneuploids show any defect in growth. One caveat of this analysis is that observed aneuploids could have adapted to handle the extra chromosomes; while this may be true in some cases, we sought common trends that might provide insights into lineage-specific effects.

Surprisingly, when strains were partitioned based on aneuploidy type, we saw no significant or major growth-rate defect in rich medium among strains that had amplified one or more chromosomes (Fig 4A). We did, however, observe a significant reduction in colony growth for strains with exclusive loss (p = 0.005, Wilcoxon test, Fig 4A). This group of 22 strains was heavily enriched for lines with higher-than-diploid base ploidy (*P* = 4×10^−8^, Fisher’s exact test), raising the possibility that reduced growth was due to ploidy burden. However, increased ploidy in otherwise euploid strains did not significantly affect growth rate compared to diploid euploids (Fig 4A). As mentioned above, strains that lost chromosomes were enriched in the Ale, African Beer, and Mixed Origin isolates, suggesting that genetic background could be a confounding feature (Fig 4B). Ale strains that had lost chromosomes were especially slow growing compared to euploids of the same clade (although these strains are also slow growing upon ploidy increases, confounding which is driving ale-strain growth defects).

**Figure 4.**
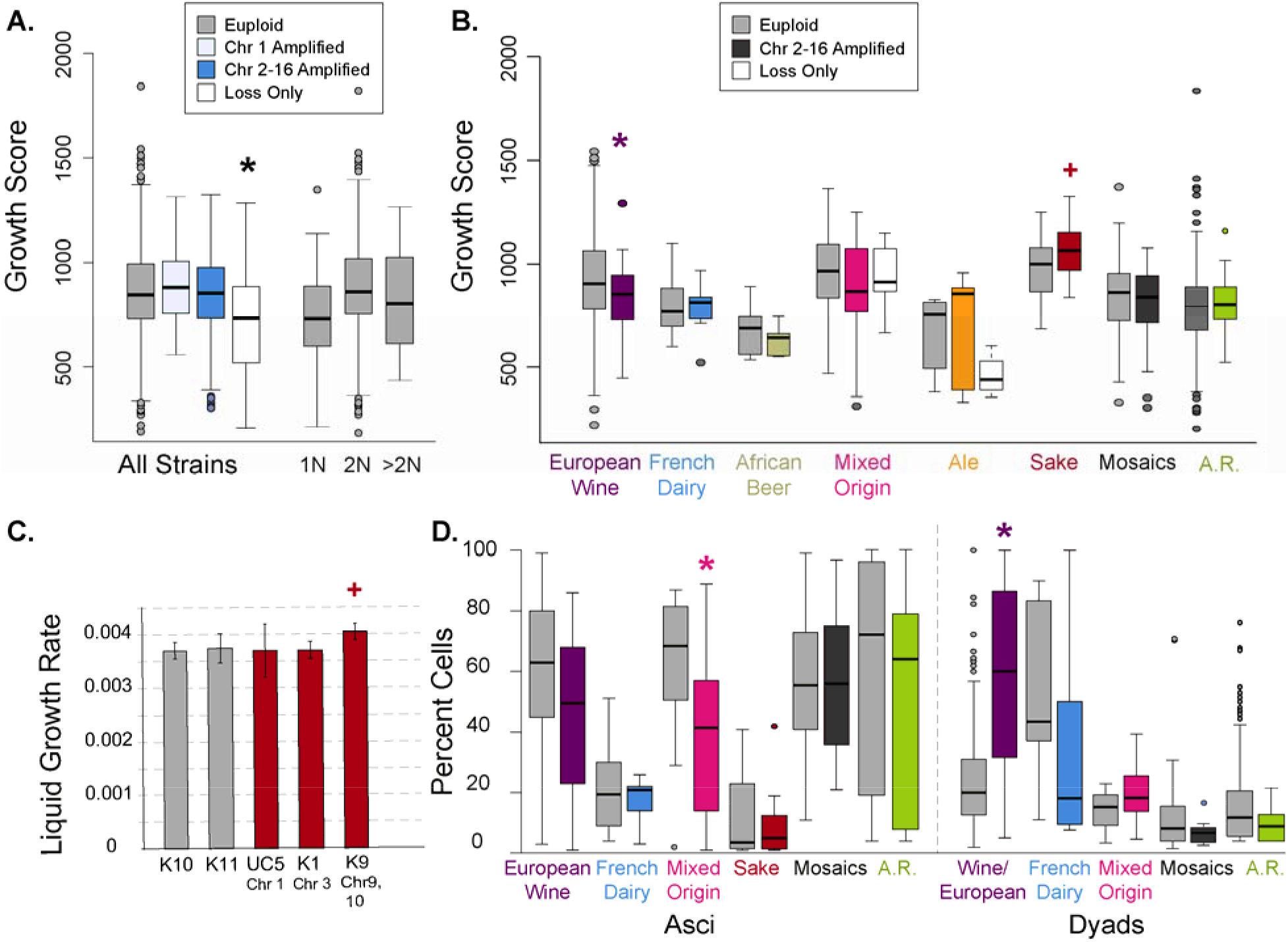
Aneuploidy presents different fitness consequences depending on strain background. A) Distribution of colony growth scores for euploid strains and strains with only Chr 1 gain, Chr 2-16 gain, or only chromosome loss according to the key. The plot also shows the distribution of fitness scores for all euploids that are haploid (1N), diploid (2N), or greater than diploid ploidy. Colored boxes represent strains with Chr 2-16 gain. B) Distribution of colony growth scores as in A but for specific clades, showing all clades with >5 aneuploids. C) Average and standard deviation of growth rates for cultures in liquid rich medium as measured in (Hose *et al.* 2015). D) Percent cells that form asci (left) or incomplete tetrads (dyads, right) after 72h in sporulation medium, for delineated clades. Groups that are statistically significantly different from the paired euploid groups are indicated with an asterisk (FDR < 0.05, Wilcoxon test) or plus (p <0.07 Wilcoxon test or T-test in C). A.R. indicates all strains excluding European Wine strains with an ancestral rate of aneuploidy (namely, those outside the French Dairy, African Beer, Mixed Origin, Ale, and Sake groups that have high rates of aneuploidy according to the logistic model).

The lack of growth defect in strains with chromosome amplification was somewhat surprising (Fig 4A). One possibility is that aggregating strains with different lineage-specifics growth rates obscures the effects of chromosome amplification. We therefore scored differences in colony growth in rich medium for individual clades that had at least five aneuploid strains (Fig 4B). Even when analyzed by clade, most groups did not show significant defects in this assay upon chromosome amplification. These included strains in the French Dairy, Mixed Origin, Ale and Sake clades, for which the aneuploid growth rates were not significantly different from euploids, considering only rich medium (Fig 4B) or all traits combined (Fig S3). In fact, sake strains with extra chromosomes displayed slightly larger colony sizes compared to euploids in that clade (*P* = 0.07, FDR = 12%). To confirm, we analyzed liquid growth rates measured for several individual strains, which grew equally well and in one case slightly better than euploid sake strains (Fig 4C (Hose *et al.* 2015)). Even when fitness scores were combined across all conditions, aneuploid sake strains showed no growth defect compared to euploid cousins (Fig S3). Although measuring liquid growth rates may reveal more subtle defects, these results reveal the absence of major defects in the strains analyzed here.

In contrast, European Wine aneuploid strains showed reduced growth rates in rich medium compared to euploids (*P* = 0.02, Wilcoxon test), a trend that persisted across many different drug conditions (FDR < 0.05, Fig S3 and 5A). Importantly, this sensitivity cannot be explained by differences in aneuploidy burden of the analyzed European Wine strains: first, there was no correlation between growth rate and change in gene content across this aneuploid group (*P* = 0.85), and second, even after removing a few outlier strains with many chromosome amplifications (see Methods), the European Wine aneuploids still fared significantly worse than euploids (*P* = 0.02, one-sided Wilcoxon test). Mosaic strains and strains from the Ale Beer group showed slightly reduced growth rates that became statistically significant when fitness traits were pooled over all conditions to boost power (FDR < 0.05, supplemental Fig S3). Interestingly, strains with ancestral rates of aneuploidy, excluding European Wine strains that scored as aneuploidy sensitive, show no statistically significant defect upon chromosome amplification, in rich media or when all conditions were combined (Fig 4 and S3). Thus, chromosome amplification has different associations with growth rate depending on genetic background, with most clades showing no major defect in sampled aneuploids.

**Figure 5.**
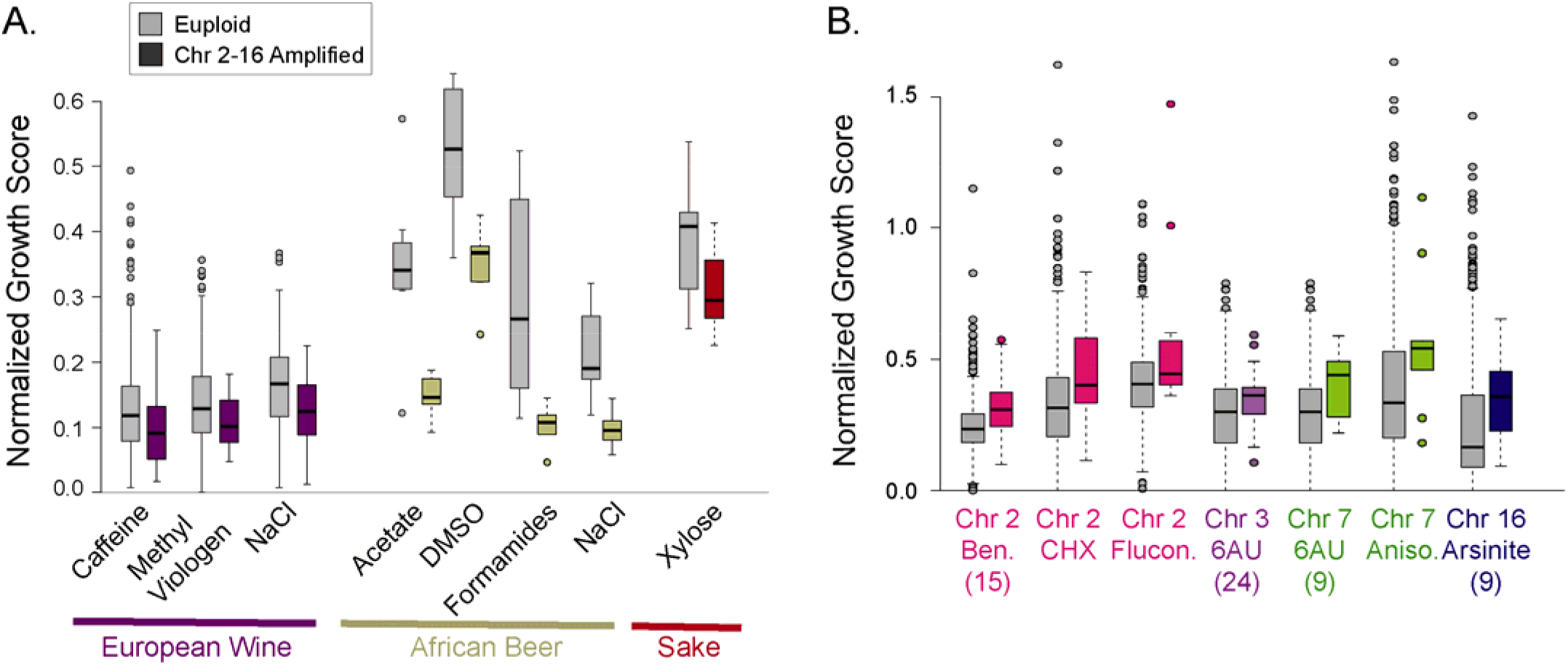
Amplification of specific chromosomes correlates with stress survival. A) Distribution of growth rates in each stress condition normalized to strain-specific growth rates in rich medium for annotated clades and stresses. In all cases shown, normalized growth of aneuploid strains was significantly worse than the comparable euploids (FDR < 0.05, Wilcoxon test). B) Distribution of growth rates as described in A for strains in which individual chromosomes were amplified. All significant results are shown in which chromosomal amplification improves relative cell growth (FDR < 0.05, Wilcoxon test).

In nature, the ability to sporulate and outcross is important both for surviving extreme periods of starvation and stress as well as to exchange genetic material. A recent study by De Chiara *et al.* characterized many of these sequenced strains for life history traits including the ability to form viable spores (asci) (De Chiara *et al.* 2020). Domesticated strains are often poor sporulators with low spore viability, proposed to result from defects segregating aneuploid chromosomes combined with a high frequency of deleterious heterozygous alleles (Gallone *et al.* 2016; Duan *et al.* 2018; De Chiara *et al.* 2020). To test if aneuploidy is associated with poor sporulation performance, we scored each clade for the propensity to form viable tetrad spores (asci) versus incomplete dyads after 72 hours in sporulation media.

Surprisingly, both the ability to sporulate and the interference associated with chromosome amplification appears clade specific. Euploid European Wine strains were among the best sporulators compared to other clades analyzed here. However, those that had amplified chromosomes bigger than Chr 1 showed a clear defect in sporulation, since most cells formed only dyads after 72 h (Fig 4D, FDR < 0.05). Defects in European Wine sporulation rate and tetrad completion (*i.e.* dyad rate) were significant after strains with outlier gene content were removed (*P*<0.01, one-sided Wilcoxon test, see Methods). Aneuploids in the Mixed Origin clade also showed a defect in asci formation compared to euploids, but no difference in dyad rate. In contrast, aneuploids in other strain groups showed no sporulation defect over euploids (FDR > 0.05). Although French Dairy and Sake strains were poor sporulators overall, extra chromosomes did not further hinder spore formation, showing that the relatively poor sporulation abilities are not caused by aneuploidy. Together, these results suggest that both the ability to sporulate and the ability to handle extra chromosomes during meiosis are influenced by clade-specific effects.

### Amplification of specific chromosomes produces passive fitness benefits

Even if having extra chromosomes is deleterious in optimal conditions, myriad studies show that aneuploidy can rapidly emerge during strong selective pressure and can present adaptive benefits (Tsai and Nelliat 2019). We found no environments in which generalized aneuploidy (*i.e.* pooled across chromosomes) was associated with improved growth; however, there were several cases where chromosome amplification exacerbated stress sensitivity, most notably for European Wine and African Beer strains grown in different environments (Fig 5A).

We next investigated phenotypic consequences of duplicating specific chromosomes independent of lineage. To control for differences in strain-specific growth rates, we normalized each strain’s stress-responsive growth to its growth in rich medium, and then assessed phenotypic gains specific to the amplification of each of the 16 yeast chromosomes. There were several important relationships (FDR < 0.05, Wilcoxon test, Fig 5B). Strains with extra Chr 16 were associated with improved growth on arsenite, strains with extra Chr 7 grew relatively better on anisomysin and 6-azauracil (the latter also associated with amplification of Chr 3), whereas the strains that amplified Chr 2 showed improved growth on several drugs. It is tempting to speculate which genes on these chromosomes contribute to the phenotypes. For example, Chr 16 carries *ARR1*, the arsenite-binding transcription factor that up-regulates defense genes including Arr3 arsenite antiporter that is also encoded on Chr 16 (Wysocki *et al.* 1997; Wysocki *et al.* 2004). Chr 7 carries two genes whose over-expression underlies tolerance to the 6-azauracil that perturbs pyrimidine biosynthesis, including pyrimidine nucleotidase *SDT1* and multidrug resistance gene *SNG1* (Shimoaraiso *et al.* 2000; Garcia-Lopez *et al.* 2010). Chr 2, which was associated with better growth on benomyl, cycloheximide, and antifungal drug fluconazole, carries the genes encoding multidrug resistance transcription factor Pdr3 and efflux pump Flr1 that gives resistance to benomyl, fluconazole, and other drugs (Broco *et al.* 1999). Verifying which genes are responsible will require specific interrogation; nonetheless, these results show that amplification of specific chromosomes can provide fitness benefits. It is interesting to note that the affected strains shown here come from very different ecologies and lineages, and thus it is unlikely that these phenotypic benefits were selected for (at least, via the same selective pressures). It instead more likely highlights the passive benefits that can arise from chromosome amplification, even in the face of a fitness cost to growth under optimal conditions in some lineages.

### Sequence variation in full-length Ssd1 does not influence aneuploidy tolerance

The extreme sensitivity of laboratory strain W303 is primarily due to a premature stop codon in the *SSD1* gene that ablates 40% of this RNA binding protein (Uesono *et al.* 1994; Uesono *et al.* 1997; Hose *et al.* 2020). Although uncommon, seven strains sequenced by Peter *et al.* reportedly carry premature stop codons (three from the *ssd1^w303^* allele), raising the possibility that extreme sensitivity to aneuploidy segregates in nature. But beyond disease alleles, it is not known if Ssd1 sequence variation contributes to aneuploidy phenotypes or frequency. We therefore analyzed Ssd1 alleles across the 1,011 strains and their association with aneuploidy phenotypes.

Nearly a third of the strains share the same Ssd1 allele, found in many vineyard strains as well as lab strain S288c. The remaining alleles differed from one another by a small number of largely private substitutions. However, both the protein tree and DNA tree revealed two major allelic classes, differentiated by two linked nonsynonymous substitutions. Allelic Class A sequence (**S**DNKQN**A** at amino acid positions 1190-1196) is ancestral in *S. paradoxus* and *S. uvarum* (Cherry 2015) and encoded in two thirds of the strains. The remaining strains carried the derived Class B polymorphisms (**G**DNKQN**P**) at those positions. Mapping Ssd1 class onto the species tree revealed that the two variants largely correlate with phylogeny (Fig 6A), with some evidence of admixture including AB heterozygotes. Interestingly, Chinese strains from primeval forest, thought to be basal to the *S. cerevisiae* species (Wang *et al.* 2012; Bing *et al.* 2014; Peter *et al.* 2018), harbor both A and B alleles as well as a hybrid A/B sequence (**G**DNKQN**A**) that likely emerged from admixture, suggesting that the B variant is anciently derived.

**Figure 6.**
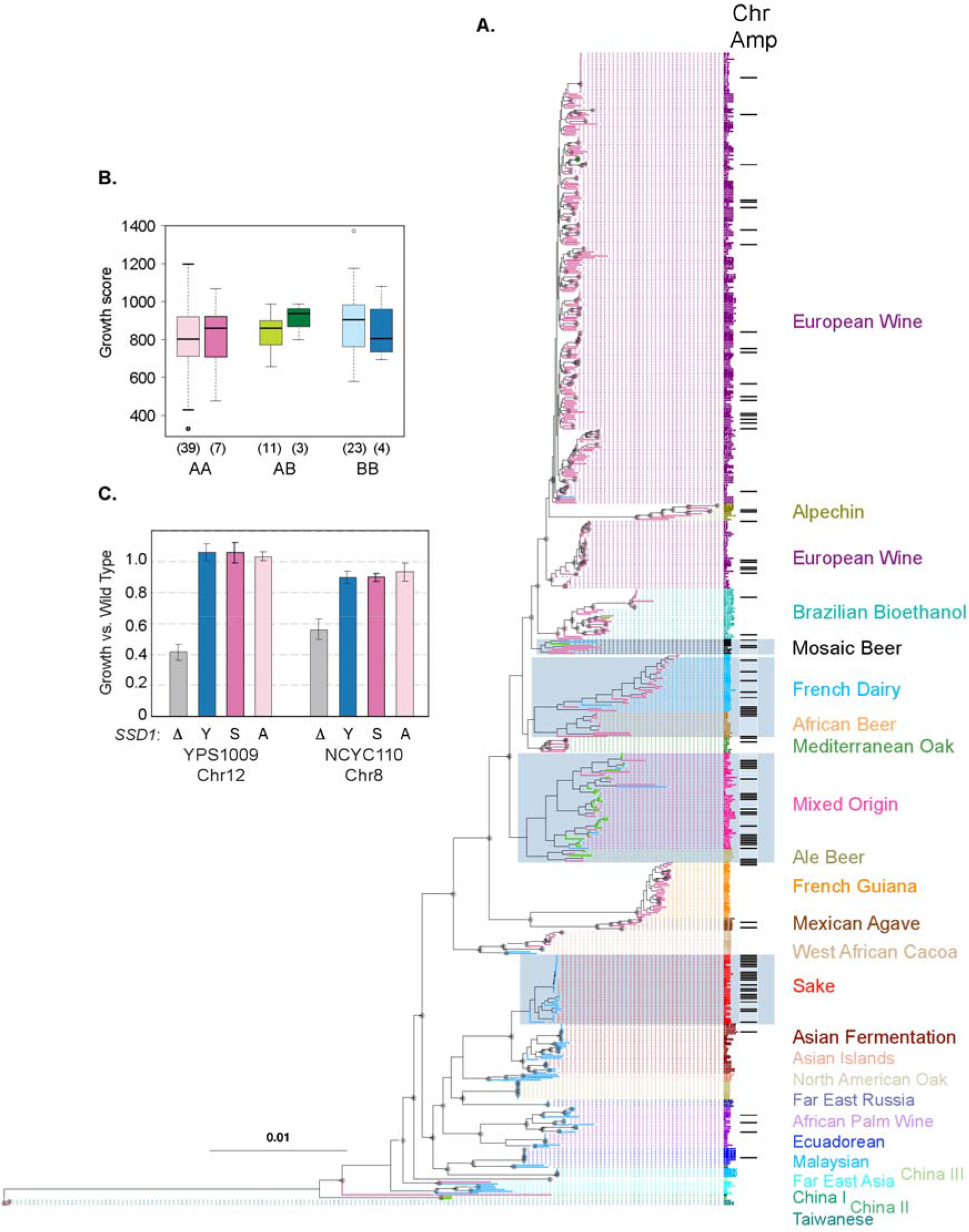
Ssd1 sequence variation does not contribute appreciably to aneuploidy tolerance. A) Maximum likelihood phylogenetic tree for the reduced set of 621 strains used for modeling (see Methods). Clades, Ssd1 allelic class (Class A, magenta edges; Class B, blue edges; A/B heterozygotes, green edges), and Chr 2-16 amplification without chromosome loss (black bars) as indicated. Strain groups with increased rates of chromosome amplification are indicated with grey boxes. Nodes with 100% bootstrap support are annotated with a grey circle. B) Colony growth scores for aneuploid *versus* euploid mosaic diploid strains expressing AA, AB, or BB Ssd1 genotypes. Number of strains per group is indicated in parentheses. No group of aneuploids was different from euploids (*P*>0.25 in all cases). C) Average and standard deviations of relative growth rate of YPS1009_Chr12 *ssd1D* cells expressing empty vector (D), the YPS1009 Ssd1 allele (“Y”, Class B), S288c Ssd1 allele (“S”, Class A), or YPS1009 Ssd1 harboring S1990, A1196 substitutions (“Y^A^”), compared to the growth rate of wild-type YPS1009_Chr12 harboring an empty vector (n=3).

Several clades enriched for chromosome amplifications express Ssd1 B variants, including the Sake clade and Mixed Origin group (which is enriched for heterozygotes); but other aneuploid groups such as the French Dairy and African Beer were largely homozygous for the AA alleles, as was the European Wine clade (Fig 6A). That both alleles were prevalent in aneuploidy-enriched clades already disfavored a generalizable role for Ssd1 allele class in aneuploidy tolerance. Nonetheless, we took two approaches to assess if allelic differences were associated with aneuploidy phenotypes. First, we reasoned that if Ssd1 protein variants contributed to differences in aneuploidy tolerance, there may be a bias in allele frequency among aneuploidy strains. However, this was not the case. There was no difference in AA, AB, and BB genotype frequencies for aneuploids with Chr 2-16 gains compared to the frequency expected based on clade proportions of the group (*P* = 0.98, Chi-squared test, see Methods). Furthermore, there was no difference in genotype frequencies in aneuploids versus euploids in the mosaic strain group (*P* = 0.2, Chi-squared test).

Second, we assessed phenotypic impacts of aneuploidy in mosaic strains with different alleles. If the Ssd1 B allele explains aneuploidy tolerance, then strains within B variants should have milder defects than strains that express A variants. Yet in all cases the mosaic aneuploids were within the euploid growth distributions, independent of Ssd1 allele class (Fig 6B). Together, these results do not support a model in which the major allele classes of Ssd1 contribute to aneuploidy tolerance.

Finally, we tested experimentally the ability of A versus B variants to complement aneuploidy tolerance in an *SSD1* deletion strain. Deletion of *SSD1* causes a major defect in the growth rate of North American oak strain YPS1009 with an extra copy of Chr 12 and in unrelated West African strain NCYC110 with a duplication of Chr 8, compared to their respective *SSD1+* wild-type aneuploid parents (Fig 6C). We found that introducing the YPS1009 B allele fully complemented S*SD1* deletion in YPS1009 and largely complemented the defect in NCYC110 (which naturally expresses a B variant). Importantly, the level of complementation was indistinguishable when S288c Class A allele was expressed, or when G1190, P1196 residues were substituted to S1190 and A1196 to mimic the A variant in a YPS1009 backbone. Thus, Ssd1 variants in the A versus B classes have no discernable effect on aneuploidy tolerance.

## DISCUSSION

Aneuploidy has played an important role in disease biology and human health as well as natural variation and evolution. An unanswered question had been the importance of genetic background in tolerating chromosomal aneuploidy, particularly the burden of chromosome amplification. Here we leveraged yeast population genomics to reflect on the forces influencing eukaryotic aneuploidy. The availability of *S. cerevisiae* genomes, phenotypes collected across environments, and strong population structure that distinguishes genetic groups provide a rich resource to address this problem. Our results shed important light on aneuploidy prevalence and tolerance in yeast that likely pertains to other organisms.

Collectively, our results reveal that genetic background alone can maximally explain variation in aneuploidy frequency and aneuploidy tolerance in *S. cerevisiae* populations. The close association between strain ecologies, ploidy, heterozygosity, and genetic background has confounded which features may contribute to differences in aneuploidy frequencies. Karyotype imbalance is more frequent as ploidy increases (Fig 1C and Storchova 2014; Zhu *et al.* 2016; Duan *et al.* 2018; Gilchrist and Stelkens 2019), consistent with models from cancer cells (Storchova and Pellman 2004), but this too is complicated in *S. cerevisiae* by the link between ploidy and clade (Fig 2). Modeling aneuploidy frequencies to decouple these factors revealed little power for ecology, ploidy, or heterozygosity beyond the explanatory power of genetic background. Our analysis predicts that increases in the propensity for chromosome duplication occurred several times in the history of the species, associated with multiple domesticated clades. While ancient admixture between the progenitors of Sake and Ale beer strains has been proposed (albeit with the ancestor of Asian fermentation strains that show low aneuploidy frequency), there is no evidence of admixture between other groups studied here (Fay *et al.* 2019), suggesting multiple genetic events in the evolution of the species. We found no evidence for selection of specific karyotypes, raising the possibility that relaxed purifying selection could have had an important role. It is interesting to note that clades that lost efficient sporulation are often enriched for aneuploid variants (Fig 4D), which could indicate that meiosis helps to purge cells with karyotype imbalance. Survival in some niches (including natural environments) likely requires meiosis and mitosis, subjecting cells to additional purifying selection compared to managed domesticated environments that do not require sex. Thus, although genetic clade best explains variation in aneuploidy rates, the result could be coupled to domestication of those lineages.

Although our models had predictive power, they explained less than 18% of the variation in aneuploidy occurrence. One possibility is that other features, including strain-specific adaptation to aneuploidy, contribute to the variance in its prevalence. But another is that aneuploidy is only slightly deleterious in the species, such that extra chromosomes appear stochastically at observable rates. The association between chromosome size/gene content and rates of chromosome imbalance (Fig 1) are consistent with a generally deleterious effect of karyotype imbalance (Gilchrist and Stelkens 2019; Tsai and Nelliat 2019). However, the effect in most strains may be far less deleterious than in the highly sensitized W303 strain. Thus, aneuploids resulting from periodic segregation defects may persist in the population for appreciable times and could enable lineages to survive stressful environments long enough to adapt through long-term mechanisms (Yona *et al.* 2012).

Clade-specific differences in the ability to handle aneuploidy stress likely further contribute to differences in aneuploidy frequency. Remarkably, most clades did not show major fitness defects upon chromosome amplification, at least based on the data and isolates studied here. This was most clear for strains in the Sake clade. Aneuploid Sake strains showed no major growth defect compared to euploid strains, based on colony sizes measured by Peter *et al.* or liquid growth rates measured in our lab for several individual strains (Hose *et al.* 2015, Fig 4). We propose that the Sake genetic lineage may be inherently tolerant of karyotype imbalance, which may explain the higher rate of aneuploidy in this clade. In fact, numerous lines of evidence suggest that these strains display unstable karyotypes. In generating meiotic products from Chr 11-amplified sake strain TCR7, Kadowaki *et al.* observed a large fraction of spores with aneuploidies not found in the parental strain, consistent with a meiotic segregation defect (Kadowaki *et al.* 2017). Another study found that some sake strains are sensitive to the microtubule poison benomyl that disrupts chromosome segregation – this sensitivity is explained by polymorphisms in Cdc55, a critical regulator of the mitotic spindle checkpoint (Goshima *et al.* 2016). Finally, anecdotal work from our lab showed variability in chromosomal duplication in sake strain K9, with different chromosomal amplifications appearing in replicate genomic analyses (Gasch *et al.* 2016). Together, these data suggest that Sake strains have unstable karyotypes due to a higher rate of segregation defects – we propose that a higher level of aneuploidy tolerance accommodates this rate. An intriguing possibility is that the higher rate of segregation errors has actually been selected for, as proposed by Kadowaki *et al.* who identified industrial benefits from chromosome amplification, regardless of which chromosome was affected (Kadowaki *et al.* 2017).

The European Wine strains provide a counterpoint to aneuploidy-tolerant strains. This clade shows low rates of aneuploidy, similar to what may be ancestral rates. However, unlike most other groups (including other strains in the ‘ancestral rate’ group defined here), aneuploid European Wine strains were slower growing than euploids in rich medium, showed sporulation defects, and were especially sensitive to stress compared to growth in optimal conditions (Figs 4–5). Growth-rate defects are unlikely to be as severe as in aneuploid W303 strains, based on past work in our lab (Hose *et al.* 2015; Gasch *et al.* 2016; Hose *et al.* 2020). Nonetheless, differences in underlying aneuploidy tolerance could have significant impacts on evolution, since aneuploidy-sensitive strains may be less likely to sample evolutionary trajectories afforded by chromosome duplication. Indeed, Filteau *et al.* previously showed that strong laboratory selective pressure produces different solutions depending on genetic background, with aneuploidy emerging in one genetic background but not another (Filteau *et al.* 2015).

An exciting and important avenue for future work is the systematic quantification of aneuploidy tolerance in the absence of selection, using lines with engineered chromosome duplications, and dissection of the genetic basis for these differences. Such defined analysis in the absence of natural selection will clarify the true impact of genetic background on variation in aneuploidy tolerance.

## Acknowledgements

We thank Joseph Schacherer for providing phenotype data described in Peter *et al.* (2018) and Mike Place and Auguste Dutcher for useful discussions and support. In memory, we thank Angelika Amon for thought-provoking and collegial discussions and for her important contributions to this field. This work was funded by NIH grant R01CA229532 to APG and NSF grant IOS 1946046 to DB and APG.

---

**Figure S1.**
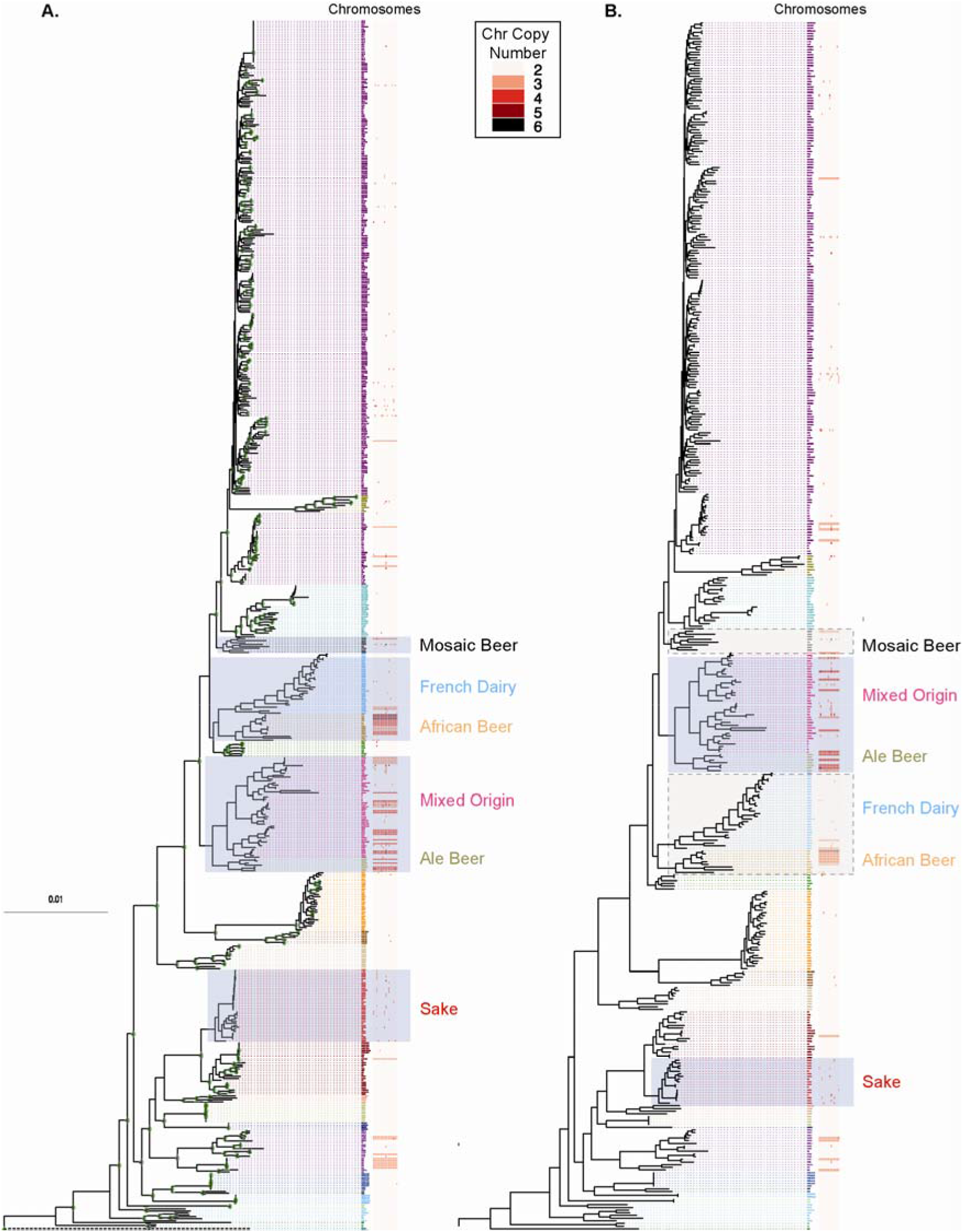
Increases in aneuploidy prevalence are not an artefact of clonal expansion of aneuploid strains. **A.** Maximum likelihood phylogenetic tree for the 621 strains used for modeling that shows the full karyotype information for each strain. See Figure 6 for all clade labels. **B.** Maximum likelihood phylogenetic tree for 453 strains (422 diploids and 31 polyploids) after removing closely related strains (genetic distance < 0.000007, see Methods). Logistic regression analysis of the 422 diploids confirmed higher rates of Chr 2-16 gain for all groups identified in the analysis of 621 strains; however, the increase was just under the significance threshold for French Dairy and African Beer (df=1, *P=*0.07) and was not significant for Mosaic Beer (df=1, *P*=0.2), likely due to reduced statistical power with fewer strains – these groups are highlighted with dashed boxes for comparison to A. Nonetheless, the model confirmed the prediction of multiple aneuploidy rate switches along the *S. cerevisiae* phylogeny. The minimal adequate model after removing close relatives showed aneuploidy rates similar to those in the larger dataset of 621 strains: an ancestral frequency of Chr 2-16 gain of 10% and elevated frequencies for Sake (P_g2-16_ = 0.54) and Ale beer and Mixed Origin clades (P_g2-16_ = 0.34).

**Figure S2.**
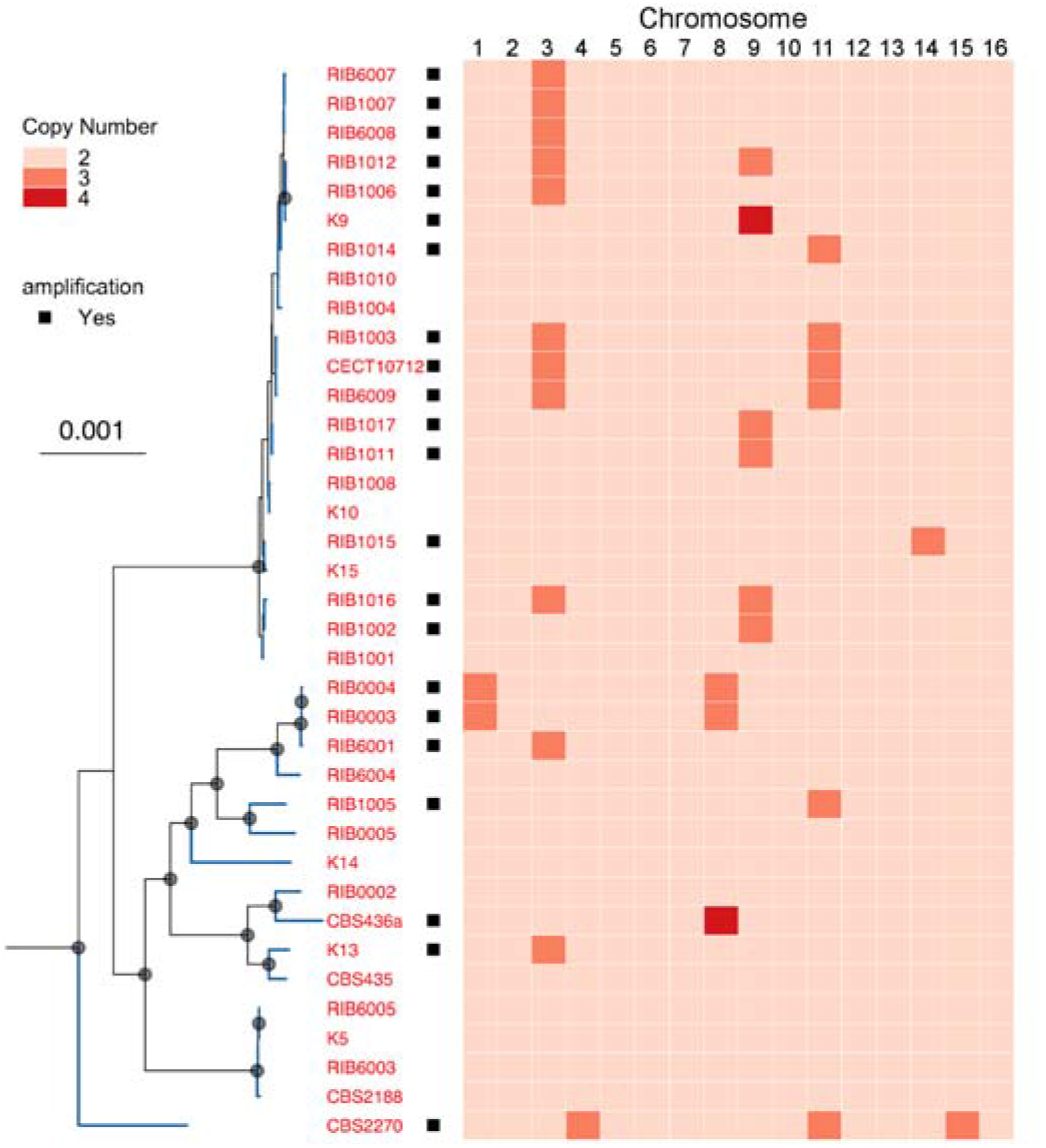
Sake karyotypes across the phylogeny. Genetic relationships among Sake strains (see also Fig S1B) compared to chromosome amplifications for each strain, according to the key. Black squares indicate aneuploid strains. Grey circles indicate nodes with 100% bootstrap support.

**Figure S3.**
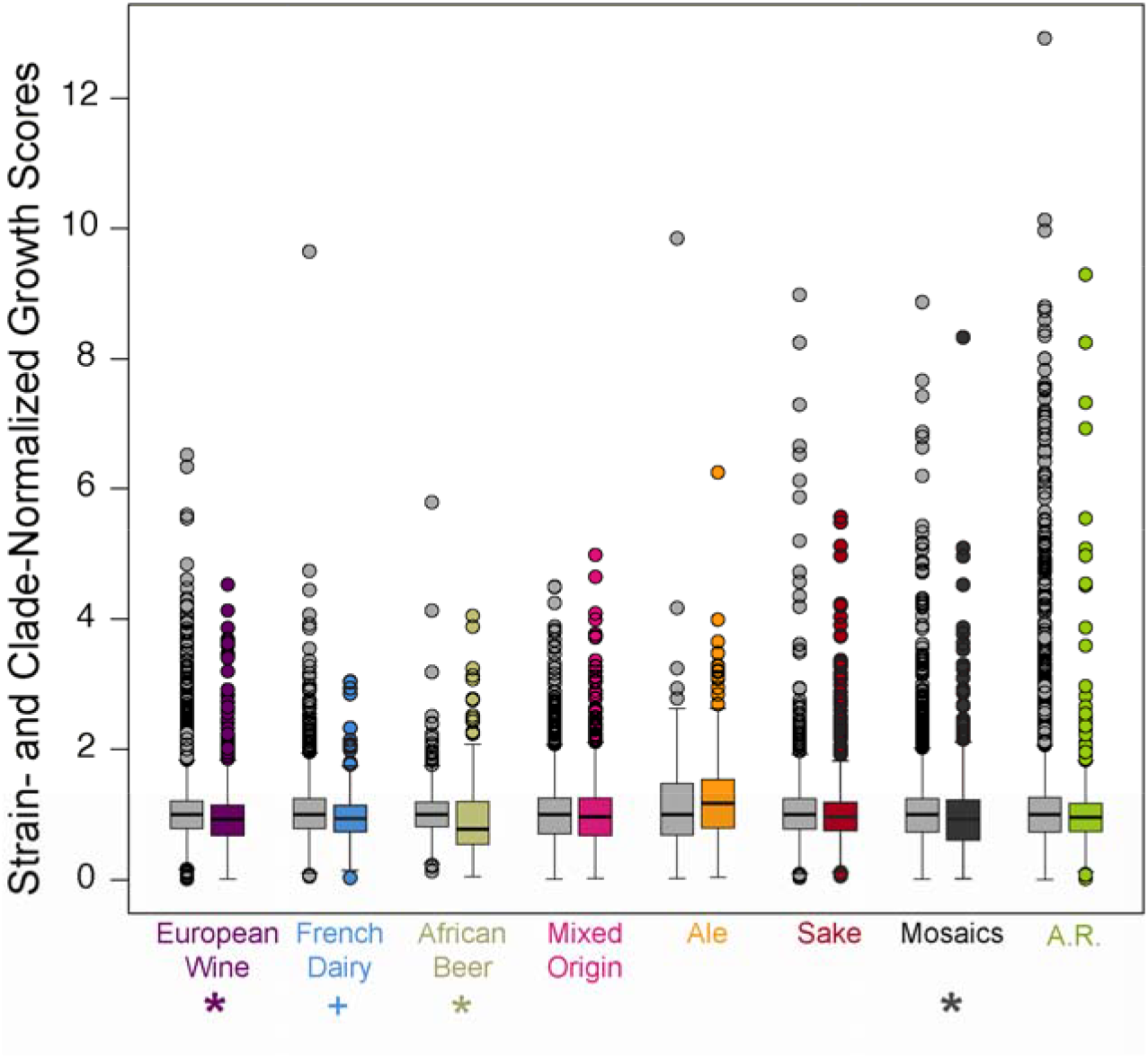
Distribution of normalized growth scores pooled across all conditions from Peter *et al.* Strain-specific growth rates measured in each of 35 conditions were normalized to each strain’s growth rate in rich medium, and then these growth scores were divided by the median score of the euploid group from that clade measured in that condition. All scores, which represent strain-, clade-, and condition-normalized growth rates were pooled and plotted, as shown in Figure 4 (asterisk, aneuploid growth defect at FDR < 0.05, + at FDR < 0.07).

